# Facility-Scale Workflows for Data Acquisition, Standardization, Machine Learning Analysis, and Reproducible Science

**DOI:** 10.64898/2026.05.06.723241

**Authors:** Sita Sirisha Madugula, Spenser R. Brown, Amber N. Bible, Ruben Millan-Solsona, Martí Checa, Lynnicia Massenburg, Alexis N. Williams, Liam Collins, Sumner B. Harris, Jennifer L. Morrell-Falvey, Scott Retterer, Rama K. Vasudevan

## Abstract

Scientific user facilities routinely generate large-scale microscopy datasets across diverse instruments and vendors, differing substantially in file formats, dimensionality, and resolution. Beyond these inconsistencies, datasets are frequently fragmented living across isolated instruments and constrained by security policies and uneven metadata practices. Consequently, tracking, standardizing, processing, and visualizing these datasets in a manner compatible with modern machine learning and autonomous experimentation workflows remains a major challenge. While existing initiatives address data archiving, standardization, or analysis individually, few provide integrated solutions that bridge instrument-level acquisition and scalable ML workflows within heterogeneous, security-constrained user facilities. Here, we establish a deployable, facility-scale infrastructure that bridges instrument-level data generation with cloud-based ML analytics while remaining compliant with institutional network constraints. Our framework integrates on-premises cloud computing, the in-house *Pycroscopy* ecosystem, and an open-source metadata management platform to transform heterogeneous microscopy datasets into standardized, ML-ready representations. We demonstrate this approach across distinct microscopy modalities through end-to-end workflows encompassing metadata capture, format harmonization, automated database ingestion, segmentation-based ML inference, and interactive visualization. By structurally separating acquisition from cloud-based analysis services, the framework enables scalable model deployment and iterative refinement without direct connectivity to instrument computers. Together, this work provides a reproducible blueprint for facility-scale data and AI infrastructure, enabling ML-ready analytics, metadata traceability, and future autonomous experimentation workflows in microscopy-driven research.

## 1. Introduction

Multidisciplinary science often requires large volumes of experimental data collected across diverse instruments, modalities, and vendors. These workflows are inherently exploratory, with evolving measurement techniques, variable data structures, and open-ended experimental design. Managing such heterogeneous datasets in a consistent, scalable, and reproducible manner remains a fundamental challenge. Conventional solutions such as Laboratory Information Management Systems (LIMS) typically excel in fixed-protocol environments^1–5^. In contrast, microscopy workflows across diverse facilities require flexible architectures that accommodate evolving measurement techniques, variable data structures, and open-ended experimental design. Such environments must support shared instrumentation and open-ended workflows operated by many users^5^. As measurement techniques evolve, data must remain interpretable and reusable beyond a single experiment or user. This requires solutions that address networking constraints, scalable storage, and metadata standardization while complying with institutional security and vendor-imposed restrictions. The rise of agentic AI further increases these requirements, calling for robust application programming interfaces (APIs) and strict standards for data accessibility, provenance, and interoperability^6^. Despite these evolving needs, the infrastructure for standardizing, processing, and visualizing data remains a fragmented landscape of ad-hoc scripts and proprietary solutions.

These gaps are widely recognized, and several large-scale initiatives have developed solutions for subsets of the problem. At the National Laboratory of the Rockies (NLR), a facility-scale infrastructure supports materials discovery by integrating data management with high-throughput computation and machine learning (ML) analysis^7^. While these systems demonstrate the value of coupling experiments with data-driven methods, they are often optimized for specific materials discovery workflows. Consequently, they lack the flexibility needed to handle high-bandwidth, heterogeneous data streams typical of multimodal microscopy in network-restricted user facilities.

Similarly, platforms such as Crucible^8^ developed at Lawrence Berkeley National Laboratory (LBNL), provides comprehensive ecosystems for centralized data acquisition, cataloging, and cloud access enabling alignment with Findable, Accessible, Interoperable, and Reusable (FAIR) principle^9–10^. However, despite robust archiving standards, a bottleneck persists at the initial data-ingestion stage. Vendor-imposed restrictions often force manual post-acquisition curation, preventing automated integration and limiting the development of workflows that generalize across imaging modalities. As a result, flexible architectures capable of real-time acquisition, processing, and metadata integration across diverse instrument platforms are still needed. Schema-driven initiatives such as NOMAD Laboratory^11–13^ and the European Open Science Cloud (EOSC)^14^, have strengthened standards for structuring and archiving heterogeneous datasets. Yet translating these standards into seamless, facility-scale FAIR implementation remains challenging, particularly at the ingestion layer.

Likewise, within the bioimaging communities, repositories like Cell Image Library, BioImage Archive, and large-scale data pipelines of Allen Institute have demonstrated the value of standardized data and rich metadata for ML-based analysis^15–17^. Although powerful, these tools are frequently decoupled from the point of acquisition. This creates a critical disconnect in environments where instrument computers are network-isolated (air-gapped), separating raw data capture from cloud-hosted analytics. Consequently, the benefits of data-sharing and reproducibility remain limited until deployable frameworks can unify the complete data lifecycle across these physical networking barriers.

To address this gap, we introduce a facility-scale deployable architecture that unifies instrument acquisition, secure data transfer, and cloud-based analysis within a continuous workflow. The framework comprises three tightly integrated components: (i) an on-premises cloud infrastructure delivering scalable storage and compute while maintaining compliance with facility security constraints, (ii) a secure bridging layer that connects air-gapped instrument systems to the cloud environment enabling automated and secure data transfer for real-time preprocessing, and (iii) a modular, instrument-agnostic software layer built around the in-house *Pycroscopy*^18-19^ package to enforce systematic metadata capture and standardization. Unlike existing data platforms that operate primarily at either the acquisition or analysis stages, our framework enables continuous, automated data flow across the full experimental lifecycle. By automatically ingesting data from air-gapped instruments into a FAIR-compliant repository, this architecture overcomes bottlenecks such as manual handling and heterogeneity barriers that hinder facility-scale integration. The architecture automatically creates an inherently ML-ready environment for real-time deployment and iterative model refinement. Our approach bridges the divide between raw experimental data and facility-scale AI, providing a scalable blueprint for future autonomous or agentic workflows.

## 2. Materials and Methods

### 2.1. System architecture

#### 2.1.1. Device enclave for instrument-cloud network isolation

The overall system is designed around a strict separation between instrument-control networks and the facility cloud environment. All analytical instruments, ranging from atomic force microscopes (AFM) to cryogenic electron microscopes (Cryo-EM), operate on isolated networks without public internet access, in compliance with facility security policies. Instrument computers operate within device enclaves that allow access to the internal cloud while prohibiting all external connectivity. This architecture ensures that instruments, regardless of modality, remain shielded from direct network exposure while still enabling data transfer into automated downstream pipelines.

#### 2.1.2. Cloud infrastructure and services

The Center for Nanophase Materials Sciences (CNMS) cloud hosts shared virtual machines (VMs) that provide compute resources for preprocessing, machine learning inference, and visualization. Cloud-accessible data shares serve as the central exchange layer between instrument datastores, analysis services, and persistent storage. All automated services, including file watchers, preprocessing pipelines, ML inference engines, and database ingestion services, operate within this cloud environment.

#### 2.1.3. Client-server orchestration layer

A client-server architecture is implemented to coordinate interactions between instrument computers, cloud VMs, file-watcher services, and the metadata database. NOMAD-Oasis^12^ is deployed to provide a searchable index of datasets and associated metadata entries. Backend services, including the NOMAD instance and orchestration components, are containerized using Docker and managed through Docker Compose. This setup enables isolated deployment of file-watcher services, preprocessing modules, and API interaction layers. Client processes running on instrument computers transmit user credentials and file paths to a central cloud VM using secure socket-based communication. Upon authentication, the server-initiated file monitoring triggers preprocessing steps and executes authenticated interactions with the NOMAD API. This architecture separates user-side acquisition systems from backend automation while maintaining secure, reproducible data handling.

### 2.2. Data acquisition and transfer

Microscopy experiments such as AFM, Cryo-EM, and optical microscopy generate raw datasets in their native instrument formats (e.g., .gwy, .ibw, .mrc, .czi). Using the instrument graphical user interface (GUI), users manually transfer acquired images to the datastore associated with the microscope or to the group managed facility storage directly. These storage locations are exposed to the cloud environment through the project-specific share drives (such as “BRaVE share” named after the project BRaVE) that are mounted onto VMs, allowing datasets to be accessed directly without requiring bulk transfer. Moreover, the shared drives also facilitate rapid transfer of datasets during or immediately after acquisition without requiring modifications to vendor-provided acquisition software. This design reflects common practice in large-scale microscopy and Cryo-EM workflows, where primary datasets remain on centralized storage and compute nodes access them through mounted network drives.

### 2.3. Automated file monitoring and preprocessing

A file-watcher script monitors the cloud data share for newly transferred files. Upon detection, the preprocessing pipeline is triggered automatically. This pipeline standardizes incoming datasets and converts them into sidpy-compatible Hierarchical Data Format (HDF5) representations^18-21^. Instrument and acquisition metadata are extracted during this conversion and stored alongside the standardized datasets. This step ensures uniform, machine-readable data representations suitable for downstream analysis and archival.

### 2.4. Model training, deployment, automated inference, and quantitative analysis via Dash interactive visualizers

Our workflow supports both off-the-shelf foundation segmentation models (e.g., CellSAM^22^, Segment Anything^23^, U-Nets^24^, YOLO^25^ and any custom-trained models by users). In this study, we employ our pre-existing YOLOv8 and YOLOv11 segmentation models fine-tuned on a manually annotated subset of AFM and Cryo-EM images of Pantoea sp. YR343 bacteria, respectively^26,27^. Trained models are deployed on CNMS cloud virtual machines (VMs), where standardized HDF5 datasets automatically trigger inference. The infrastructure allows models to be retrained, updated, and executed directly within the cloud environment, enabling iterative refinement without requiring instrument-level access. Segmentation masks generated during inference are stored alongside the original datasets and associated metadata within the cloud environment. Backend processing pipelines compute quantitative bacterial properties, including cell count, surface area, eccentricity, and morphology-derived descriptors. Results are delivered through integrated Dash-based visualization interfaces for interactive inspection and analysis.

### 2.5. NOMAD-Oasis deployment

An internal NOMAD-Oasis instance is deployed on the CNMS on-premises cloud infrastructure to provide metadata management, dataset indexing, and search capabilities. NOMAD exposes REST APIs for authentication, data upload, querying, and publishing, along with a web-based interface for interactive access. User authentication is performed through the NOMAD API, where credentials are exchanged for time-limited access tokens. These tokens are used for all subsequent API operations that are managed programmatically by a client-server orchestration layer that coordinates file monitoring and ingestion services. A graphical client application running on instrument computers collects folder paths and NOMAD authentication details and transmits them securely to a cloud-hosted server through SSL-protected sockets. The server validates credentials, launches automated file-watching and preprocessing services. Once initiated, the “watcher” constantly monitors shared storage for new datasets triggering automatic HDF5 format conversion and ingestion workflow. Dataset registration and metadata indexing are subsequently performed within the NOMAD-Oasis API.

### 2.6. Custom schema and QR code generation

In addition to the instrument- or acquisition-specific metadata that are automatically ingested with standardized HDF5 files, we implemented a mechanism to capture additional experimental metadata records and associate them with physical samples. For this purpose, a custom electronic laboratory notebook schema, BRaVESampleELN, is developed using YAML-based specifications. The schema captures entries for sample-specific information such as sample identifiers, preparation conditions, biological or chemical context and imaging annotations. Each dataset ingested into NOMAD is assigned a globally unique upload identifier (upload_ID) serving as a reference to a single ingested sample. For each upload_ID, metadata files related to the sample are uploaded as separate entries and every entry receives a unique entry_ID. This serves as a persistent identifier linking the sample and its associated metadata, which can be accessed and visualized through the NOMAD graphical interface as a complete metadata record.

To link these digital records with physical samples, a QR code-based system is implemented that generates a QR code for each metadata entry. A python-based utility queries the NOMAD instance for entries within an upload_ID, constructs the corresponding NOMAD GUI URL by combining the base interface address with the upload_ID and entry_ID, which are encoded by QR codes. Scanning the QR code opens the complete metadata record of a sample directly within the NOMAD interface. To facilitate practical laboratory use, a mapping file associating entry_ID values with human-defined sample names is also generated. This mapping preserves readability for users while maintaining compatibility with NOMAD identifiers. QR codes produced through this workflow can be stored digitally or printed and attached to physical samples, directly, enabling rapid retrieval of metadata during analysis. This setup enables granular tracking of individual samples, datasets, and associated metadata across experimental workflows.

## 3. Results and Discussion

### 3.1. Secure data exchange enables automated facility-scale data ingestion

The enclave-based architecture described in Section 2.1.1 enabled instrument computers operating on isolated networks to exchange data with the CNMS cloud environment while operating within institutional network-isolated environments. Using this framework, representative datasets from heterogeneous microscopy platforms, including both Windows- and Linux-based acquisition systems, are successfully processed via the automated ingestion and analysis pipeline while remaining compliant with facility security requirements. To validate the architecture, we accessed AFM and cryo-EM datasets via the data share, allowing file-watching and preprocessing services to operate directly on this drive without requiring bulk transfers. This facilitated data standardization, metadata capture, and programmatic upload of datasets into NOMAD-Oasis for further indexing and referencing while keeping primary data stationed on facility storage. Although demonstrated on representative instruments, the architecture is designed to extend to facility-level storage, requiring a mere reconfiguration of mount points. This configuration establishes a flexible framework for automated preprocessing, metadata capture, and downstream analysis across multiple instruments and modalities

### 3.2. Dataset standardization using *Pycroscopy* and Sidpy

To enable consistent downstream processing, incoming heterogenous datasets are standardized using the *pycroscopy* ecosystem^28^, which is designed for structured handling of multidimensional image datasets. At the core of this ecosystem is the sidpy^29^ dataset abstraction, providing a unified representation of scientific datasets coupling numerical data arrays with structured metadata describing dimensions, physical quantities, units, and experimental context. Raw datasets from instruments are read using modality-specific readers and converted into sidpy datasets, where their spatial, spectral, and temporal dimensions are explicitly defined and annotated. This explicit encoding of data semantics distinguishes sidpy datasets from conventional image stacks or vendor-specific file formats, enabling consistent handling of complex, high-dimensional microscopy data across modalities. The standardized datasets are then serialized into hierarchical data format (HDF5) files which serve as the primary on-disk representation for downstream workflows. HDF5 provides a scalable, self-describing file format capable of efficiently storing large microscopy datasets together with their associated metadata.

Importantly, the internal file structure within HDF5 is organized to remain compatible with the NeXus26 data standard. This ensures interoperability with a broad range of external analysis and visualization tools while preserving the richer metadata representation provided by sidpy. Embedding metadata directly within the dataset ensures that standardized files are self-contained and portable, reducing the risk of metadata loss and simplifying ingestion into databases such as NOMAD-Oasis. Therefore, *Pycroscopy* and sidpy are used in the standardization layer, providing a practical bridge to transform raw heterogeneous datasets from multiple instruments into a uniform, generalizable across imaging modalities representation that is immediately amenable to automated preprocessing, visualization, and ML-based analysis.

### 3.3. Facility-scale dataset and metadata management via NOMAD-Oasis

The local NOMAD-Oasis instance deployed on CNMS cloud environment, provides a facility-wide platform for metadata and dataset indexing and search. Harmonized HDF5-formatted datasets are ingested into NOMAD-Oasis via its API, where structured schemas organize metadata and experimental context. While primary datasets remain on facility storage, NOMAD maintains searchable records and archival representations supporting rapid discovery of datasets based upon their experimental parameters, samples, and associated attributes in future. This deployment consolidates facility knowledge by providing a centralized index of experiments and metadata, enabling datasets to be located, compared, and reanalyzed over time without requiring changes to the underlying storage infrastructure (Figure 1).

**Figure 1.**
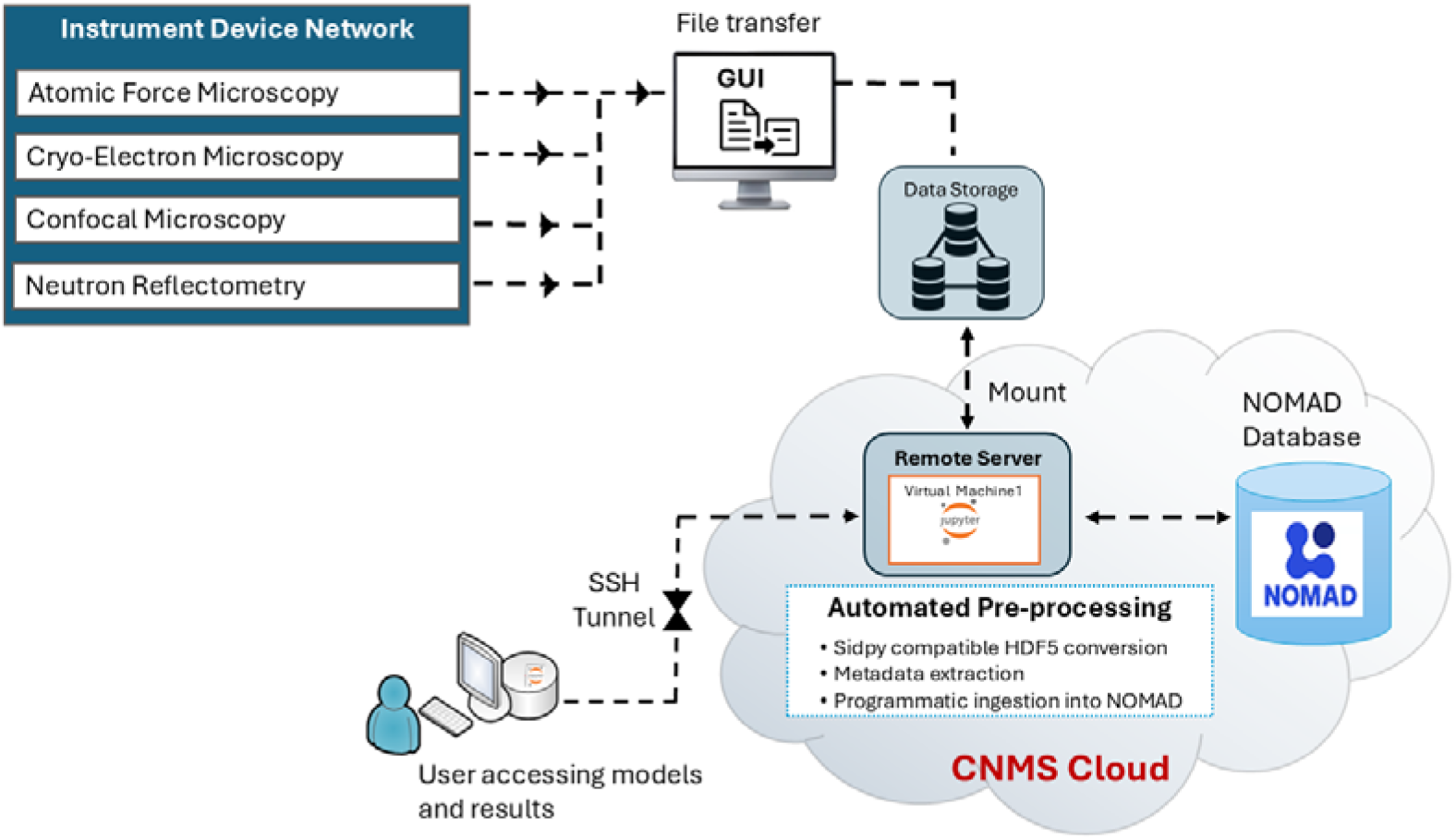
Facility scale end-to-end microscopy data orchestration and metadata management pipeline. The workflow illustrates the integration of instrument networks with the cloud infrastructure for automated pre-processing and FAIR-compliant data management. Raw datasets are converted to HDF5 format and indexed within NOMAD-Oasis, separating physical storage from searchable metadata to support rapid experimental discovery and data provenance.

We validated our design using representative multimodal datasets such as .mrc, .gwy, .ibw, and .czi files originating from various microscopy instruments mounted via the shared drive. The raw datasets are quickly converted to standardized HDF5 files, which are automatically detected by the file-watcher and ingested into NOMAD-Oasis without any manual intervention. Within NOMAD, each dataset receives a unique upload_ID, enabling traceable registration and retrieval. NOMAD also indexed metadata allowing datasets to be searched and accessed through the web interface immediately after preprocessing. The automated pipeline enabled datasets to transition from acquisition and preprocessing to indexed, searchable records, facilitating downstream ML analysis. While the current implementation uses NOMAD for metadata management, the framework is designed to be database-agnostic and allows substitution with alternative federated database management systems such as DataFed^29^, which is currently being integrated for services such as sample tracking via QR code, metadata indexing and provenance tracking of data with metadata. Together, this architecture separates data standardization and metadata management, allowing flexible, user-friendly capture workflows to coexist with facility-scale data governance. This separation is essential for supporting large, heterogeneous user facilities where both human usability and infrastructure scalability must be addressed simultaneously.

### 3.4. Custom schema enabled sample lifecycle tracking

Standardized datasets preserve acquisition metadata extracted from instrument files, but experimental context such as sample preparation records, instrument logbook entries, or other laboratory annotations must typically be recorded manually. To capture such contextual metadata, we implemented the custom metadata schemas discussed in Section 2.7, demonstrated here through sample-level lifecycle tracking. The architecture supports incremental, multi-user metadata entry, allowing different users to contribute information at successive stages of a sample’s lifecycle. Here we demonstrate sample-level metadata ingestion via *BRaVESampleELN* using representative confocal microscopy (.czi) datasets. Information such as sample identifiers, preparation details, experimental conditions, and imaging annotations are captured systematically and stored as structured records. The schema also captures details of sample handling and experimental progression, such as transfer events, imaging modality, and the user responsible for the next analysis step in the experiment allowing stage-wise recording of sample history. These records are further tracked by aligning them with metadata entries from subsequent experiments performed using different imaging modalities. Metadata entries from these subsequent experiments are ingested as separate entries associated with the same sample, allowing records from different experimental stages to be viewed within the NOMAD environment. This approach demonstrates a method for incremental metadata capture enabling tracking of sample history across experiments and instruments. Work is ongoing to embed automatically extracted metadata and ELN records alongside standardized datasets to strengthen the linkage between experimental data and laboratory context.

### 3.5. Linking physical samples to digital metadata records via QR codes

In many multidisciplinary scientific infrastructures, physical samples and their digital records are often managed separately, making it difficult to quickly retrieve experimental context. Our QR-code-based association mechanism links physical samples with their respective metadata records in NOMAD-Oasis. We demonstrate this on confocal microscopy datasets, although the same procedure is also applied for AFM and Cryo-EM datasets. Raw proprietary image datasets generated from Zeiss microscope (.czi), once standardized, are ingested into NOMAD-Oasis, supplemented with sample-level metadata, and rendered via the NOMAD URL which is encoded by the QR code. Scanning the code retrieves the complete metadata record of the sample including information like sample preparation details, experimental conditions, imaging annotations, and records of subsequent analyses performed on the sample. The URL not only displays file-level metadata but also links each record to its Upload_ID assigned during dataset ingestion, ensuring traceability to the original upload. This enables retrieval of the complete experimental context of a sample rather than just its name or identifier (Figure 2). Additionally, the mapping files associate human-readable names of the samples with their NOMAD Entry_IDs, simplifying sample searching, and cross-referencing while preserving the identifiers for future database operations. QR codes produced through this workflow for bacterial samples used in confocal imaging are saved digitally. However, this setup allows users to print QR codes and attach them directly to physical samples, enabling rapid access to metadata during handling, imaging, and analysis. These QR codes can be scanned using standard tools or approved devices, providing the flexibility needed to retrieve metadata across diverse facility environments. This setup represents a systematic method to link physical samples with standardized datasets, and metadata, enabling practical tracking of samples and their associated experimental context.

**Figure 2.**
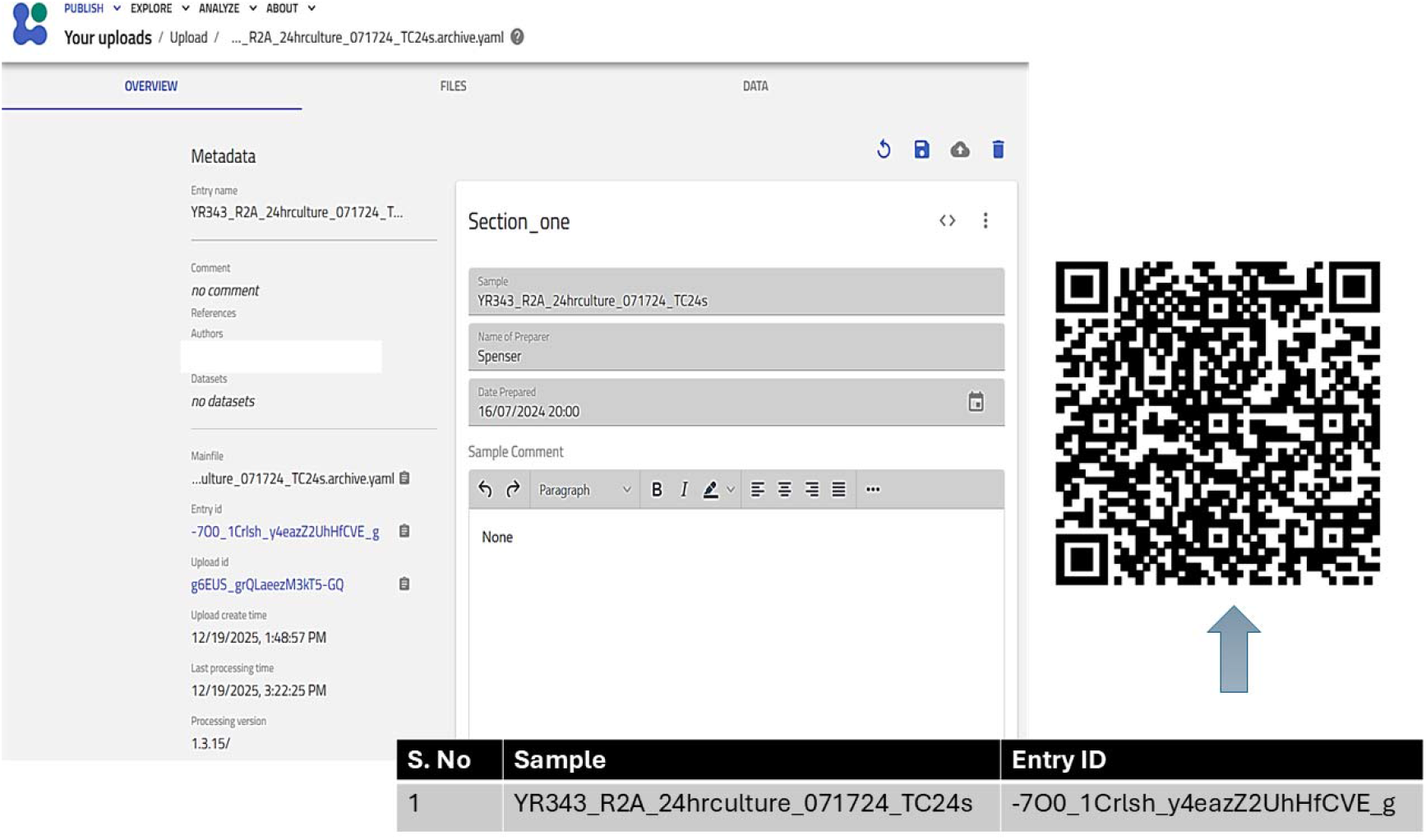
Sample metadata displayed via custom schema and tracked using QR codes. The figure demonstrates a QR-code-based association mechanism used to link physical samples to their digital entries. This setup uses custom schemas within NOMAD-Oasis to render sample-level metadata (left). The resulting digital record is linked to the physical sample by encoding its unique NOMAD URL into a QR code (right). Scanning the code resolves to the NOMAD URL, providing immediate access to the sample’s complete experimental context, including preparation details, imaging annotations, and assigned Entry_ID and Upload_ID.

### 3.6. End-to-end ML-enabled facility workflow and deployment

The integrated infrastructure described above enables dynamic deployment of ML-based analysis within a secure facility environment. Standardized datasets, automated preprocessing, and structured metadata management together provide the foundation for scalable, closed-loop microscopy workflows. As a representative demonstration, we applied the complete pipeline to AFM and Cryo-EM datasets, implementing automated image segmentation and quantitative analysis within the CNMS cloud infrastructure. Harmonized HDF5-formatted datasets generated during preprocessing are immediately compatible with cloud-hosted segmentation models, quantitative analysis routines, and interactive Dash-based visualization interfaces. Figure 3(a-b) illustrates the inspection of segmentation outputs and derived quantitative measurements for Cryo-EM and AFM datasets, respectively. These components collectively establish a closed-loop system linking acquisition, standardization, ML inference, visualization, and metadata indexing without direct interaction with instrument-control systems.

**Figure 3.**
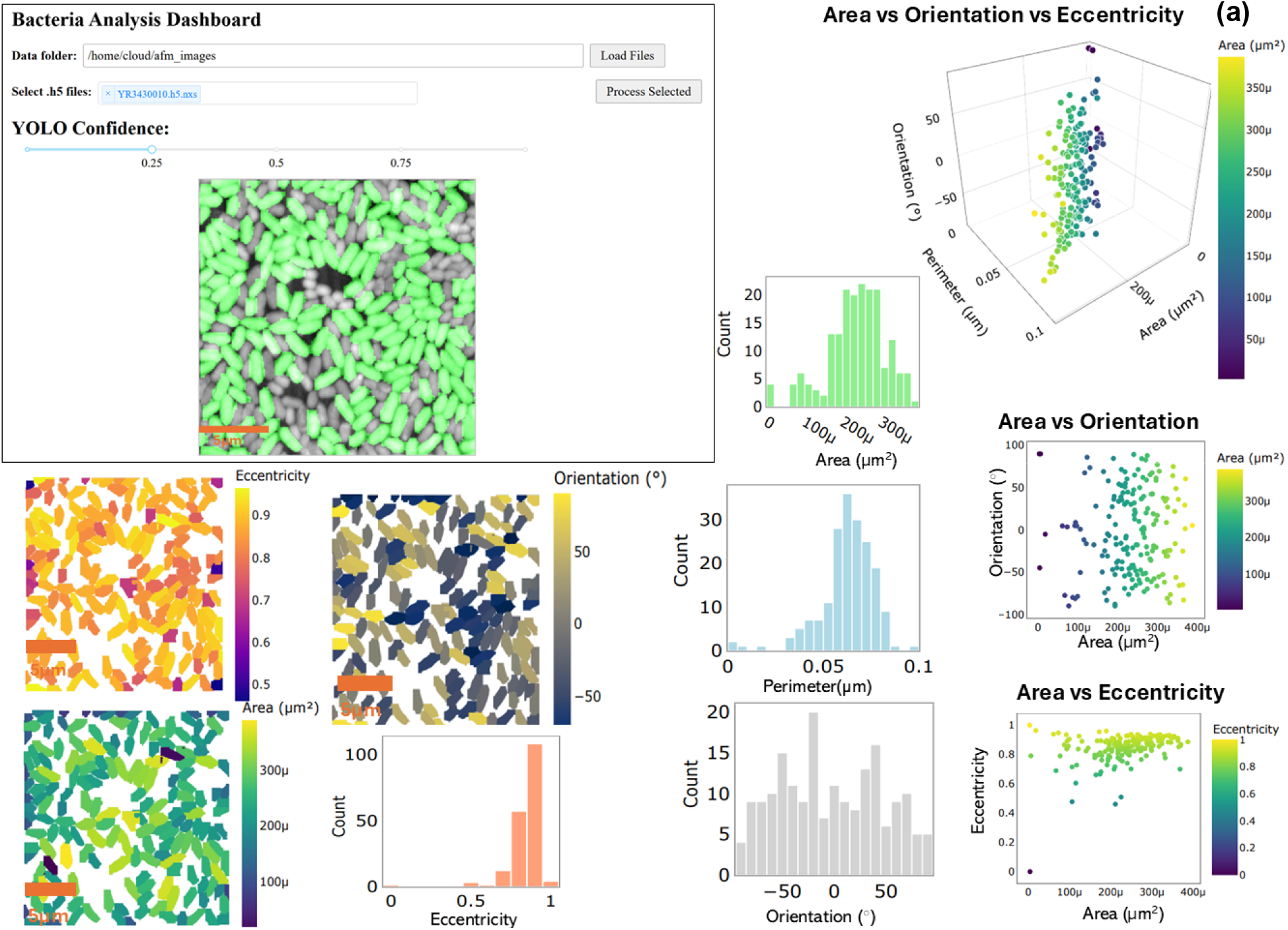
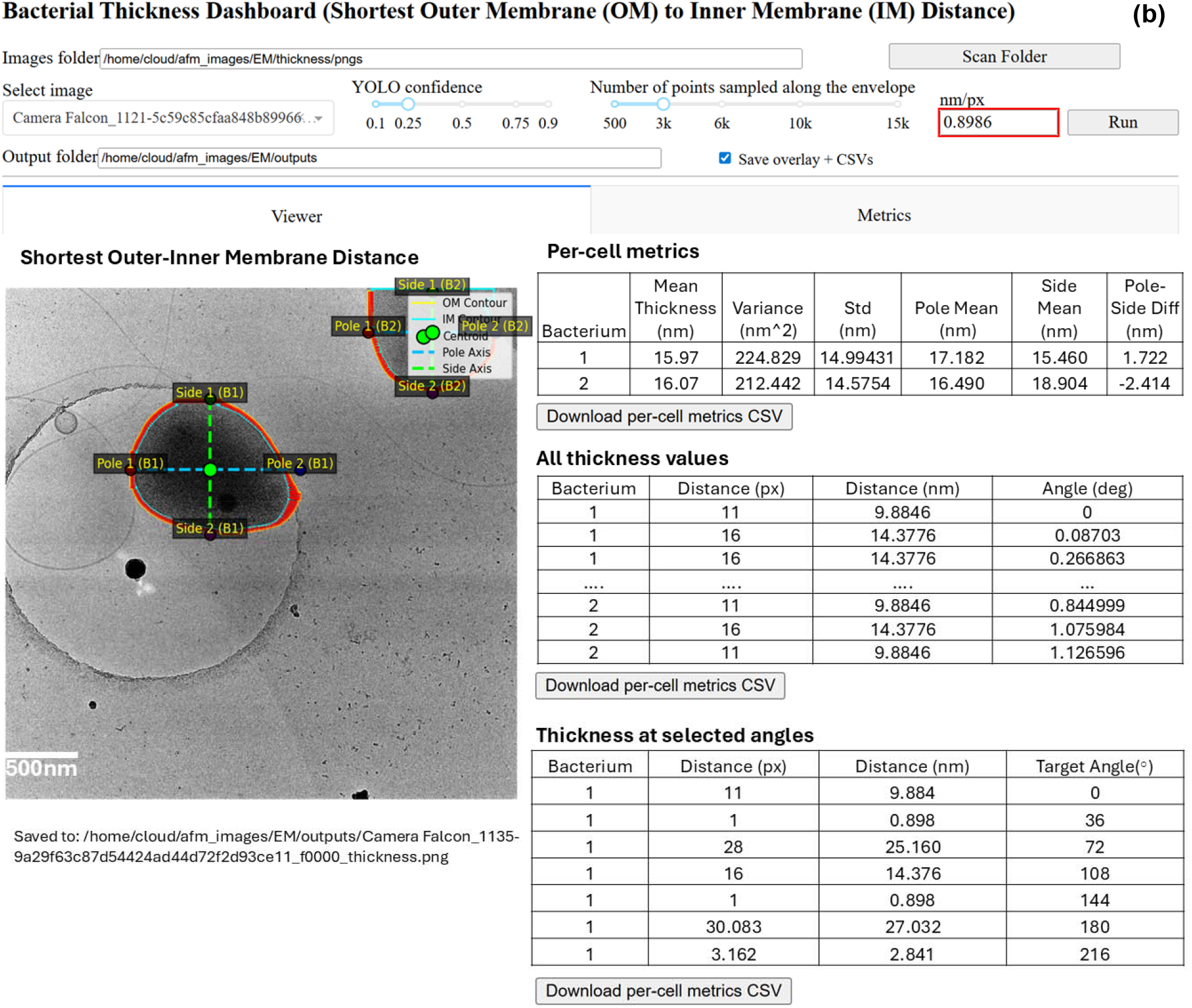
Dash-based interactive visualization interface used to inspect segmentation results and the derived quantitative measurements for (a) AFM and (b) Cryo-EM datasets respectively. The dashboards provide visual platforms to bridge the gap between raw ML outputs and scientific interpretation. By overlaying masks directly onto raw images, it allows users to verify the fidelity of segmentations before proceeding with analysis. The interface displays complex datasets into multidimensional plots and tabular summaries, providing a visual means to compare various quantitative measurements simultaneously. In AFM datasets, this includes morphological properties like area, orientation, and eccentricity. Similarly, for Cryo-EM datasets, the dashboard enables the visualization of cell envelop masks providing high-precision structural metrics, such as membrane thickness recorded as detailed tabulations. This centralized approach allows users to identify population trends and outliers across different imaging modalities while interactively exploring how model parameters influence the resulting biological insights.

Figure 4 presents the unified end-to-end workflow demonstrated on AFM datasets. Raw datasets acquired using AFM are generated in native instrument formats and transferred to the facility data store via a GUI and then to the project specific data share. These files are thereafter transferred to CNMS cloud enabling automated transfer of image files without exposing instrument computers to external networks.

**Figure 4.**
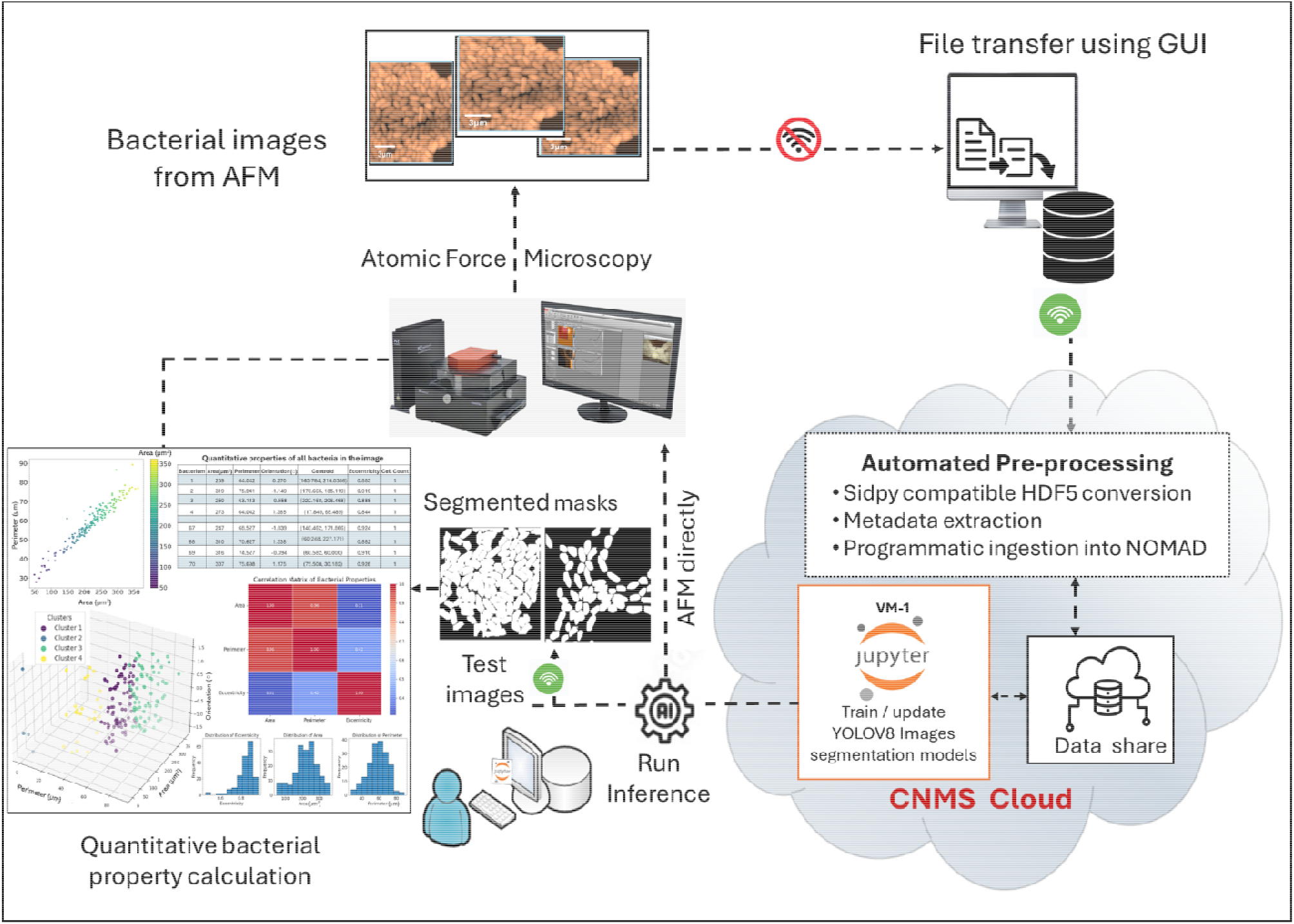
End-to-end facility workflow for ML-enabled analysis of AFM bacterial imaging data across instrument and cloud networks. Raw AFM images acquired in native formats are transferred from instrument-side systems to a local datastore via a graphical user interface, providing a secure handoff between the instrument network and the facility cloud. Data is automatically transferred to the CNMS cloud, where standardized preprocessing, HDF5 conversion, metadata extraction, and ingestion into NOMAD-Oasis are performed. ML-based segmentation models placed in the cloud are deployed on the newly ingested datasets for inference and the resulting segmentation masks are used for quantitative bacterial property calculations and interactive visualization.

Within the CNMS cloud, incoming datasets undergo an automated preprocessing pipeline that includes data standardization, conversion into HDF5 formats, and registration of a corresponding data entry in the local NOMAD-Oasis instance. During preprocessing, the instrument-specific metadata is automatically pushed along with HDF5 files, ensuring consistent capture of acquisition parameters without manual intervention. Further, users dynamically enter any additional sample-level metadata are associated with the datasets using the custom or inbuilt schemas of NOMAD-Oasis depending on the nature of the metadata. Harmonized HDF5-formatted datasets are then made available for downstream ML analysis within the cloud infrastructure. In this demonstration, we employ our pre-existing YOLOv8s-based model trained on manually annotated AFM images^24^.

Newly standardized datasets automatically trigger inference. Alternatively, models can be dynamically trained, updated and executed directly on the CNMS VMs as new data becomes available. The resulting segmentation masks pass through the ML-based analysis pipeline that quantifies bacterial morphological and spatial descriptors. Outputs are delivered through Dash-based visualization tools, enabling interactive validation without manual file transfer. Where required, derived results could be routed back to instrument-side environments to support additional experimental interpretation. The versatility of this standardized architecture is further demonstrated by its successful deployment on Cryo-EM datasets. While the imaging modality differs, the underlying workflow remains identical. Using our pre-existing YOLOv11-seg^21^ and U-Net segmentation models, we achieve high-fidelity automated analysis of Cryo-EM datasets with the same efficiency as that seen in the AFM pipeline.

Importantly, the workflow presented here is not specific to AFM or image segmentation. By decoupling data acquisition, standardization, metadata management, and analysis execution, our architecture provides a general-purpose framework for deploying diverse data-driven and machine learning analyses across diverse instrument platforms in a shared facility environment. Automated data transfer between instruments and cloud networks maintains institutional network isolation while enabling scalable cloud-based analysis. Throughout this workflow, data, metadata and analysis outputs remain co-located within the facility infrastructure, with harmonized HDF5-based representations serving as the unifying layer that enables reproducible, interoperable workflows across instruments, models, and analysis tools.

### 3.7. Human-in-the-loop model refinement and incremental training

To demonstrate model adaptability, we implemented a human-in-the-loop (HIL) refinement over the existing YOLOv8-seg model described earlier^26^. Segmentation masks predicted for incoming AFM images using existing models are first inspected to identify missing bacterial predictions or inaccuracies where the model fails to identify all bacteria within the images accurately. In our experiment, the initial model is specifically optimized for AFM images of *Pantoea sp. YR343* bacteria, and the model requires precise calibration to the spatial scaling and morphological proportions. This requires retraining the existing model to improve its ability to detect all bacteria accurately. To achieve this, we integrated SimuScan^30^, a synthetic data generator that allows for targeted model refinement through user-guided parameter corrections specified via the package’s configuration file. By using expert manual measurements of bacteria from representative original images, we recalibrate SimuScan’s parameters to generate synthetic training datasets.

These recalibrated parameters are then used to generate additional synthetic training images that capture the bacterial morphology. The resulting datasets are further incorporated into an incremental fine-tuning workflow, allowing the existing segmentation model to be updated (fine-tuned) using newly generated data, without requiring additional manual annotation or retraining from scratch. The resulting fine-tuned model is saved as a new model together with the associated training configuration and synthetic dataset parameters, allowing the refinement step to be documented and reproduced. These updated models can be reused for subsequent analysis within the workflow. This iterative process allows the segmentation model to progressively adapt to variations in sample morphology, or experimental protocols encountered over time. In our experiment, the effectiveness of HIL intervention is evidenced by the substantial improvement in detection performance of the updated model compared to the base model. Following the targeted recalibration, evaluation metrics improved across mAP50 (0.94 from 0.73), Precision (0.99 from 0.73), and Recall (0.98 from 0.78), while the number of high-confidence bacterial predictions (>0.8 confidence) increased to 255 (from 108) in a representative test set image. This transition demonstrates the workflow’s efficacy in adapting to specific imaging morphologies through human-guided incremental training without the need for exhaustive manual annotation. In our implementation, the human-in-the-loop component consists of expert inspection of predicted segmentation masks to identify bacteria missed by the existing model. More generally, the framework allows human intervention when model outputs indicate potential failure cases or when adjustments are required for specific experimental conditions. In other implementations, such interventions could also be triggered through automated uncertainty detection or similar system-level prompts.

## 4. Conclusions

We present an end-to-end, facility-scale framework that integrates automated data ingestion, incremental metadata capture, cloud-based ML, and interactive visualization within a secure and reproducible environment. By separating instrument operation from downstream preprocessing and analysis while preserving standardized data formats and persistent metadata, the workflow enables scalable, near real-time computation under realistic security constraints.

More importantly, this framework changes how multi-disciplinary science can collect and refine datasets. Data is no longer static record processed after acquisition. They become continuously structured, searchable, and computation-ready assets from the moment they are generated. Although demonstrated using AFM or Cryo-EM bacterial imaging and segmentation models, the architecture is modality- and model-agnostic. It provides a general foundation for embedding data-driven and ML-enabled analysis directly into routine facility workflows. This infrastructure establishes a practical pathway toward ML-native scientific user facilities, where data acquisition, analysis, and iterative model improvement can occur continuously within secure institutional environments, enabling future autonomous and data-driven experimentation.

## Acknowledgements

This work was supported by the U.S. Department of Energy, Office of Science FWP ERKCZ64, Structure Guided Design of Materials to Optimize the Abiotic-Biotic Material Interface, as part of the Biopreparedness Research Virtual Environment (BRaVE) initiative. AFM, EM imaging was conducted as part of a user project at the Center for Nanophase Materials Sciences (CNMS), which is a US Department of Energy, Office of Science User Facility at Oak Ridge National Laboratory. SSM and RKV thank Sarthak Kapoor, Hampus Näsström and the FAIRmat team for their support in setting up NOMAD-Oasis and schema development.

## Author contributions

S.S.M. developed the methodology, ML pipelines, result analysis, code repository and manuscript writing, while S.B.H. contributed to developing DL models for Cryo-EM image analysis. S.R.B., A.N.B., J.L.M.-F. contributed to growing the bacterial cultures. R.M.S., M.C., L.C. carried out experimental AFM imaging while M.L. and A.N.W. acquired the Cryo-EM images. R.K.V. conceptualized the methodology and supervised the overall study. The original draft was written by S.S.M. while all authors reviewed and edited the manuscript. Funding was acquired by S.R.

## Data Availability

The datasets, models and all architectural scripts are available at https://github.com/Sireesiru/AI-ML-Computational-workflows.

## Conflicts of Interest

All the authors declare no conflict of interest.

## Notes

### Competing Interest Statement

The authors have declared no competing interest.

